# High-resolution postmortem MRI of the human infant brain: cortical reconstruction and anatomical parcellation

**DOI:** 10.64898/2026.05.07.722301

**Authors:** Pulkit Khandelwal, Sala Young, Nathan Xi Ngo, Paul A. Yushkevich, Andre van der Kouwe, Robin L. Haynes, Hannah C. Kinney, Lilla Zöllei

## Abstract

High-resolution postmortem magnetic resonance imaging enables detailed examination of brain anatomy at spatial scales not achievable *in vivo* and provides a unique opportunity to link morphometric measurements with the underlying pathology. Despite these advantages, robust computational tools for automated anatomical segmentation and cortical surface reconstruction remain limited, particularly in postmortem infant brains. Incomplete myelination, thinner cortical ribbons, small-scale neuroanatomy, evolving tissue contrast, fixation-induced signal alterations, and variability in postmortem preparation make standard neuroimaging pipelines unsuitable for postmortem infant MRI. In this work, we introduce a unique high-resolution multisequence postmortem infant MRI dataset and a unified computational framework that combines deep learning-based volumetric segmentation with surface-based cortical reconstruction and anatomical parcellation in native subject-space resolution. The framework is designed to generalize across diverse postmortem MRI acquisition protocols, spatial resolutions, tissue preparation conditions, and specimen characteristics while remaining robust to substantial variability in image contrast, tissue deformation, fixation-induced intensity changes, background signal characteristics, and anatomical variability encountered in postmortem infant MRI. We benchmark our framework against widely used contrast-agnostic and foundational brain segmentation models, demonstrating improved anatomical consistency and segmentation performance across heterogeneous high-resolution postmortem infant datasets. Our method enables morphometric analysis in native postmortem space, providing the same downstream quantitative analyses routinely available for *in vivo* developmental neuroimaging. The complete framework is released as open-source software with command-line workflows, containers, and comprehensive documentation to facilitate reproducible postmortem infant MRI analysis as part of the `purple-mri` package.

## 1 Introduction

Magnetic resonance imaging (MRI) is central to developmental neuroscience, enabling quantitative characterization of early cortical gray matter (GM) maturation, white matter (WM) organization, and emerging structural networks during infancy [1,2]. Automated segmentation, surface reconstruction, and atlas-based parcellation [3,4] translate *in vivo* MRI into region-specific morphometric measures supporting studies of normative development and early vulnerability [5]. However, *in vivo* infant MRI is constrained by motion, limited acquisition time, and safety considerations, restricting spatial resolution and contrast diversity [6]. Consequently, fine-scale cortical folding, laminar organization, and small subcortical structures remain difficult to resolve.

Postmortem MRI provides a complementary approach. By eliminating motion and permitting prolonged acquisitions, postmortem imaging enables substantially higher spatial resolution and field strengths than feasible *in vivo* [7,8,9]. In infancy, this permits detailed assessment of cortical ribbon geometry, deep gray nuclei, and brainstem–forebrain circuitry [10]. Such datasets are particularly relevant in neuropathological contexts, including sudden infant death syndrome where subtle structural alterations are hypothesized but challenging to detect *in vivo* [11].

In adult neuroimaging, high-resolution postmortem MRI has linked morphometric signatures directly to histopathology in neurodegenerative diseases [12], revealing microstructural correlates of pathology [13,14]. Standard neuroimaging pipelines, both classical (e.g. FreeSurfer [15], FSL [16]) and deep learning–based (e.g. FastSurfer [17]), are not well suited to the analysis of postmortem MRI due to distinct imaging characteristics [18], motivating the development of postmortem–specific frameworks. For example, purple-mri [19,20] enables native-space cortical reconstruction and atlas-based parcellation for high-resolution postmortem data, while NextBrain [21,22] leverages a probabilistic histology-derived atlas within a Bayesian segmentation framework for cortical and subcortical regions in both *in vivo* and postmortem MRI.

Synthetic data–driven deep learning frameworks [23], including SynthSeg [24] and SuperSynth [25], achieve contrast-, scanner-, and resolution-agnostic segmentation via large-scale domain randomization. However, they were developed for adult *in vivo* MRI and assume mature cortical geometry and stable gray–white matter contrast, limiting generalization to infant anatomy characterized by thinner cortical ribbons, ongoing myelination, evolving tissue contrast, and smaller subcortical structures. Infant-specific synthetic approaches such as Infant-Synthseg [26] and BabySeg [27] model developmental contrast trajectories in *in vivo* imaging, addressing limitations of traditional infant pipelines (e.g. InfantFreeSurfer [28], iBEAT [29]). Nevertheless, these methods remain grounded in *in vivo* signal assumptions and do not model fixation-induced relaxation shifts, laminar cortical heterogeneity, immersion-media background signal, or specimen-specific artifacts characteristic of high-resolution postmortem infant MRI. Thus, there is a lack of a framework which addresses the combined challenges of postmortem acquisition, developing infant neuroanatomy, and high-resolution multisequence imaging.

### Contributions

In this work, we first present a unique multi-sequence postmortem infant MRI dataset acquired at sub-millimeter resolution. Second, we introduce a unified segmentation-to-surface computational framework tailored to postmortem infant MRI that integrates deep learning-based volumetric tissue segmentation, topology correction, and native-space cortical reconstruction with atlas-based parcellation. This enables robust downstream cortical morphometry, including region-wise volumetric analysis and vertex-wise cortical thickness estimation. Third, we show that the framework performs consistently across the image sequences, acquisition variations, and specimen characteristics while remaining robust to altered tissue contrast, fixation-induced intensity changes, background signal, deformation, and local tissue damage. Finally, we conduct comprehensive benchmarking against five established foundation-model and contrast-agnostic segmentation approaches, demonstrating improved robustness and anatomical consistency in high-resolution postmortem infant imaging.

## 2 Materials and Methods

### 2.1 Dataset

We acquired a unique postmortem infant brain MRI dataset (12 specimens; 8 male, 4 female), scanned on a 3 T Siemens scanner at 0.5*×*0.5*×*0.5 mm^3^ resolution using multiple sequences: multi-echo fast low-angle shot (FLASH), diffusion MRI (dMRI), and true fast imaging with steady-state free precession (TRUFI). Multi-echo FLASH acquisitions used variable flip angles (10–50^*°*^), polarity variations, and both short and long echo times to enhance contrast diversity and sensitivity to tissue relaxation. dMRI was acquired with high-angular resolution (80–90 directions; 10–15 *b*0 volumes). TRUFI sequences employed 8 phase-cycling angles (0^*°*^–315^*°*^), with select specimens additionally scanned at multiple flip angles (e.g., 30^*°*^, 70^*°*^) to improve signal uniformity and boundary delineation. Thereafter, quantitative T1 [30] and PD [31] maps were derived from multi-echo FLASH using [15], fitting a multi-parameter FLASH model across flip angles (Figs. 2, 3).

**Fig 1.**
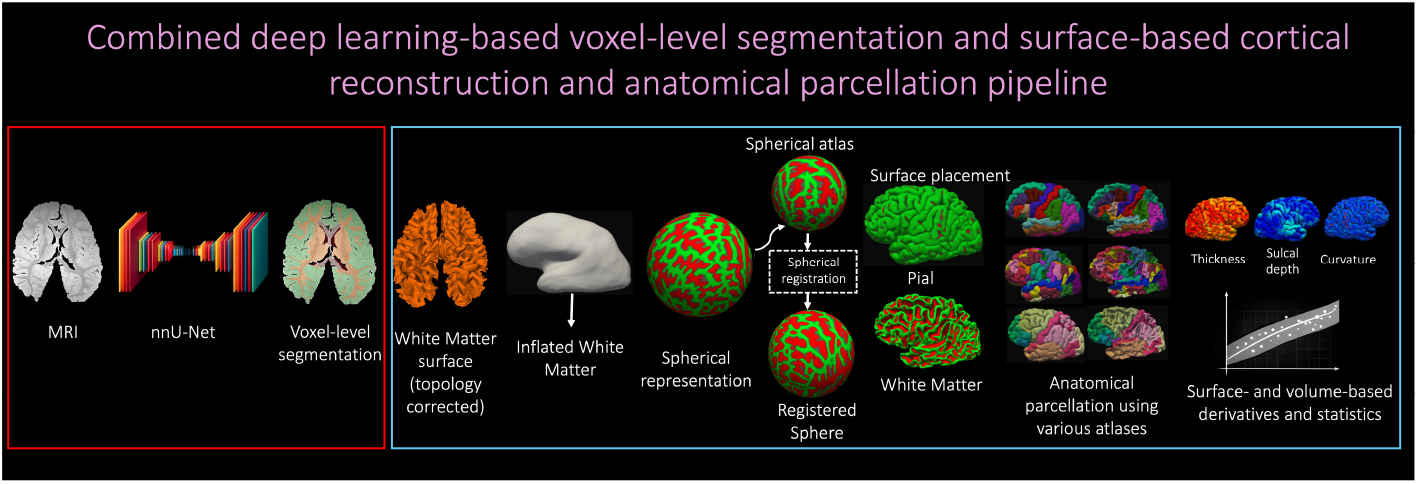
Deep learning-based segmentation (left) and surface-based cortical reconstruction and anatomical parcellation (right) pipeline as explained in Section 2.2.

**Fig 2.**
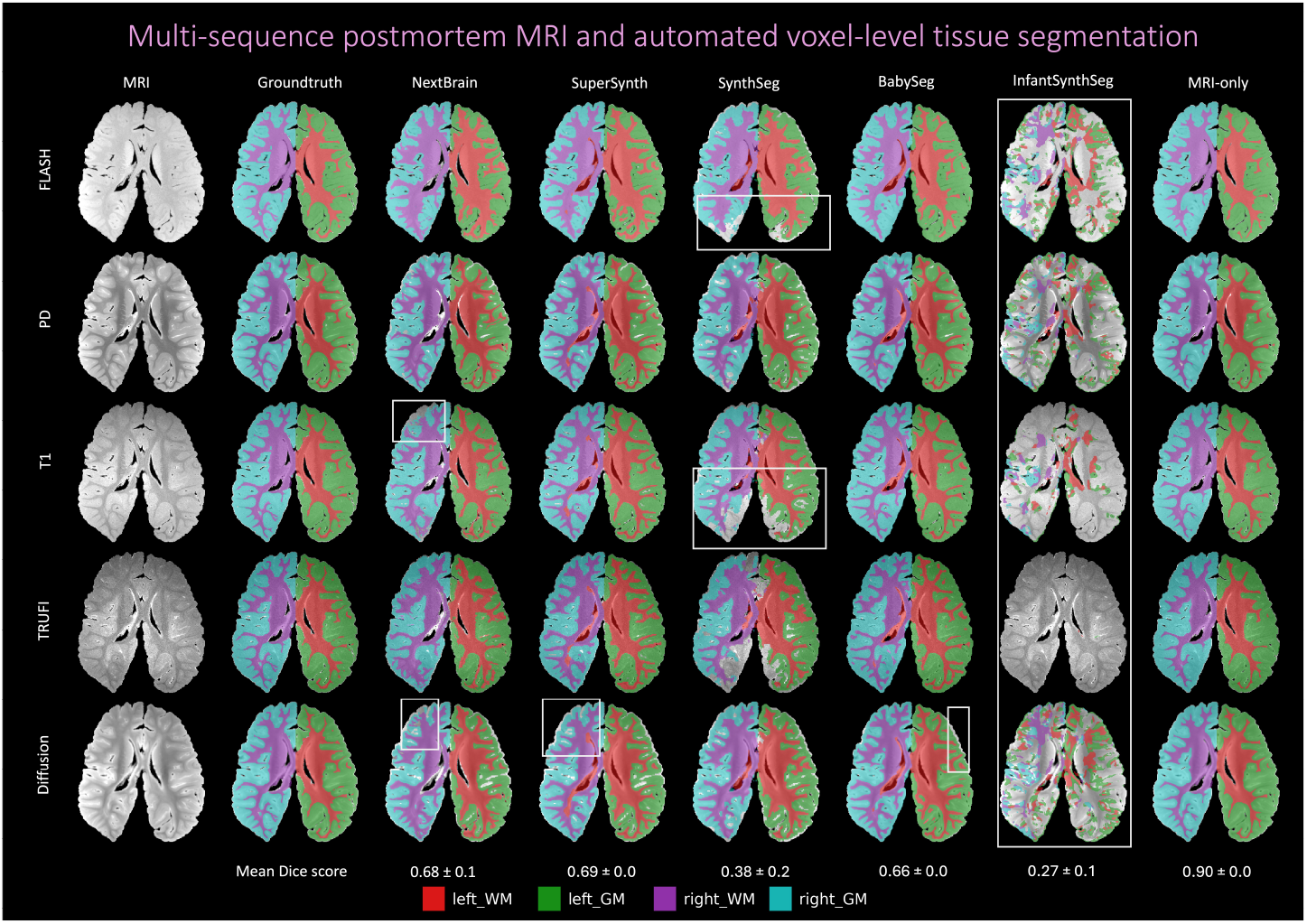
Multi-sequence postmortem human infant MRI. Comparison of various voxel-level deep learning-based tissue segmentations (Section 3.1) for a single specimen.

**Fig 3.**
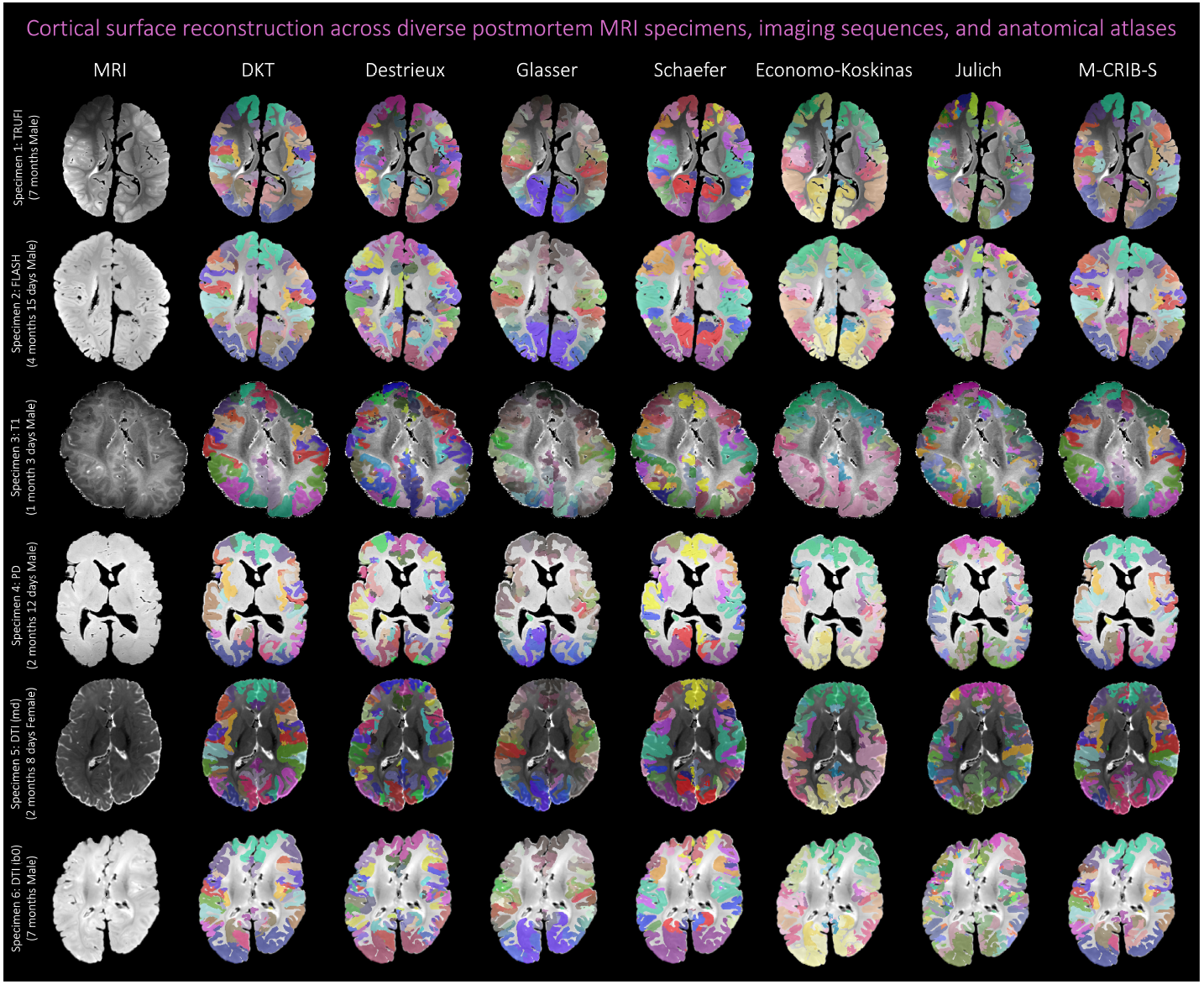
Postmortem human infant MRI. Anatomical parcellation of different specimens using a variety of atlases on various sequences.

The mean age at death was 3.1*±*2.8 months (range: 1.0–10 months), and the mean postmortem interval (PMI) was 21.8*±*13.1 hours (range: 3.5–47 hours); all specimens were fixed in formalin for at least 30 days before imaging. A trained research assistant manually generated anatomical labels for cortical GM, WM, and hippocampus (minimum set of anatomical labels required for downstream surface-reconstruction pipeline) on the FLASH scans, which provided superior tissue contrast for manual delineation. For each specimen, all other imaging sequences were rigidly aligned [32] to the FLASH space, and labels defined in FLASH space were then also used as ground truth for all other four sequences.

### 2.2 Tissue segmentation and anatomical parcellation framework

Inspired by [19,15], we introduce a framework for infant postmortem MRI that integrates deep learning-based volumetric tissue segmentation (Fig. 1 red box) with topology correction and classical surface-based reconstruction, inflation, registration, and anatomical parcellation (Fig. 1 blue box).

#### Deep learning-based volumetric tissue segmentation

Given an input structural volume *I*(**x**), a 3D network *f*_*θ*_ predicts voxel-wise tissue labels Ŝ(**x**) *∈ {*1, … , *L}* , including cortical GM, WM, and hippocampus.

#### Surface-based cortical reconstruction and anatomical parcellation

The refined segmentation 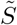 drives surface reconstruction following [19,15,33,34]. A volume is formed by treating WM and hippocampus as foreground and cortical GM as background, then tessellated to generate an initial WM surface *S*_WM_, which is topologically corrected [35]. The surface is inflated to a spherical representation *S*_sphere_ and registered via curvature-driven spherical alignment to the Buckner40 atlas space [36]. The pial surface *S*_pial_ is obtained by outward evolution of *S*_WM_ under smoothness and boundary attraction constraints. For *in vivo* MRI, evolution is guided by ∥ ∇*I*(**x**) ∥ to detect the GM–CSF interface. However, in high-resolution postmortem infant MRI, multi-echo contrast induces substantial intra-cortical intensity heterogeneity, leading to premature termination within GM and cortical thickness underestimation. To address this, surface evolution is constrained by segmentation-derived anatomical boundaries the surface is restricted to the GM support of 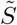 and terminated at the segmentation-defined GM-background interface. This stabilizes pial placement, prevents sulcal fusion, and yields anatomically consistent thickness estimates. Finally, labels from multiple atlases are transferred from spherical atlas space to the subject-specific spherical representation.

## 3. Experiments and Results

### 3.1 Experiments

The framework was evaluated against foundation segmentation models using PyTorch [37] on an NVIDIA A5000 GPU.

#### Methods

For the voxel-based segmentation pipeline, we used nnU-Net [38] as the 3D backbone because of its robustness across heterogeneous imaging domains and competitive performance [39]. We trained an MRI-only nnU-Net model using real postmortem infant data and benchmarked it against segmentation methods: NextBrain, SuperSynth, SynthSeg, InfantSynthSeg, and BabySeg. All methods were evaluated using their recommended preprocessing and inference settings.

#### Training details and evaluation

The dataset comprised 12 specimens, each contributing five input volumes: FLASH, dMRI, TRUFI, and derived quantitative T1 and proton-density maps, for a total of 60 volumes. MRI-based segmentation experiments used five-fold cross-validation across specimens. The nnU-Net model was optimized using a combined Dice and cross-entropy loss, and segmentation accuracy was assessed against expert-guided labels using the Dice similarity coefficient. Results were reported per structure as mean*±*SD, aggregated across sequences, hemispheres, and folds.

Sequence-specific predictions were fused by majority voting to generate a consensus segmentation for each specimen, which was then used for cortical surface reconstruction and atlas-based parcellation. Desikan-Killiany-Tourville (DKT) atlas-based regional gray matter volumes and cortical thickness measurements were extracted from the postmortem parcellations. These measurements were descriptively compared with *in vivo* developmental trajectories spanning the same 0–10-month age range, derived using Infant FreeSurfer [28] from 155 participants in the Baby Connectome Project [40], as an external anatomical plausibility reference.

## Results

### Voxel-based tissue segmentation

Table 1 highlights a clear domain and imaging hierarchy. The MRI-only model, trained directly on real postmortem infant data, achieved the highest performance (0.90*±*0.0), demonstrating the importance of domain-specific supervision for high-resolution postmortem infant MRI. Adult *in vivo* foundation models showed partial transfer but remained limited by domain and anatomical shift: SuperSynth (0.69 *±* 0.0) and NextBrain (0.68*±*0.1) achieved the strongest performance among comparison methods, whereas SynthSeg (0.38*±* 0.2) degraded markedly. Infant-focused models did not consistently improve performance, with BabySeg achieving moderate transfer (0.66*±*0.0) and InfantSynthSeg performing poorly (0.27*±*0.1), underscoring that infant anatomy alone is insufficient without postmortem domain alignment. Together, these results show that training directly on real postmortem infant MRI provides substantially more robust and anatomically consistent tissue segmentation than existing contrast-agnostic or foundation segmentation models. Fig. 2 qualitatively illustrates these differences, highlighting improved cortical and hippocampal delineation by the nnU-Net model.

**Table 1.** Dice similarity coefficients (↑mean ± std. dev.) averaged across left and right hemispheres. Also shown are the training strategies’ summary for each model: anatomy (adult vs infant), domain (*in vivo* vs *ex vivo*), imaging (real vs synthetic).

| Method | Anatomy | Imaging characteristics | WM<sub>avg</sub> | GM<sub>avg</sub> | HIPPO<sub>avg</sub> | Overall |
| --- | --- | --- | --- | --- | --- | --- |
| NextBrain | *in vivo*-adult | Bayesian atlas | 0.70 ± 0.1 | 0.77 ± 0.1 | 0.58 ± 0.2 | 0.68 ± 0.1 |
| SuperSynth | *in vivo*-adult | *in vivo*-syn | 0.71 ± 0.1 | 0.79 ± 0.1 | 0.58 ± 0.1 | 0.69 ± 0.0 |
| SynthSeg | *in vivo*-adult | *in vivo*-syn | 0.41 ± 0.3 | 0.55 ± 0.2 | 0.17 ± 0.3 | 0.38 ± 0.2 |
| BabySeg | *in vivo*-infant | *in vivo*-syn | 0.69 ± 0.1 | 0.79 ± 0.1 | 0.50 ± 0.1 | 0.66 ± 0.0 |
| InfantSynthSeg | *in vivo*-infant | *in vivo*-syn | 0.35 ± 0.2 | 0.40 ± 0.2 | 0.08 ± 0.2 | 0.27 ± 0.1 |
| MRI-only (oracle) | *ex vivo*-infant | *ex vivo*-real | 0.87 ± 0.0 | 0.92 ± 0.0 | 0.92 ± 0.0 | **0.90 ± 0.0** |

### Surface-based cortical reconstruction and parcellation

Using the MRI-only model, sequence-specific segmentations were fused by majority voting to generate a consensus label map for each specimen, which served as input to the surface reconstruction and parcellation pipeline. Full cortical reconstruction, surface inflation, and volumetric atlas projection required approximately 45 minutes per specimen. The pipeline generated multiple anatomical parcellations, including M-CRIB-S [41], DKT [42], Schaefer [43], Julich [44], Glasser [45], and von Economo–Koskinas [46]. The inclusion of infant-specific template-based parcellations such as M-CRIB-S further supports anatomically appropriate labeling of the developing cortex. The bottom row shows reconstructed cortical and white matter surfaces for the first specimen using the M-CRIB-S atlas. Figs. 3 and 4 demonstrate robust cortical reconstruction and atlas-based parcellation across multiple postmortem infant specimens and MRI sequences. Anatomically consistent regional boundaries are maintained despite low gray-white matter contrast, pronounced anterior-posterior intensity gradients, fixation-induced signal heterogeneity, tissue deformation, and localized imaging artifacts. For quantitative evaluation, Desikan-Killiany-Tourville (DKT) atlas-based regional gray matter volumes and cortical thickness measurements were extracted from the postmortem parcellations and compared with normative 0-10 month *in vivo* developmental trajectories derived from the Baby Connectome Project (Fig. 5). Postmortem morphometric measurements followed expected developmental trends and remained within physiologically plausible ranges, supporting the anatomical fidelity of both volumetric segmentation and downstream surface-based modeling.

**Fig 4.**
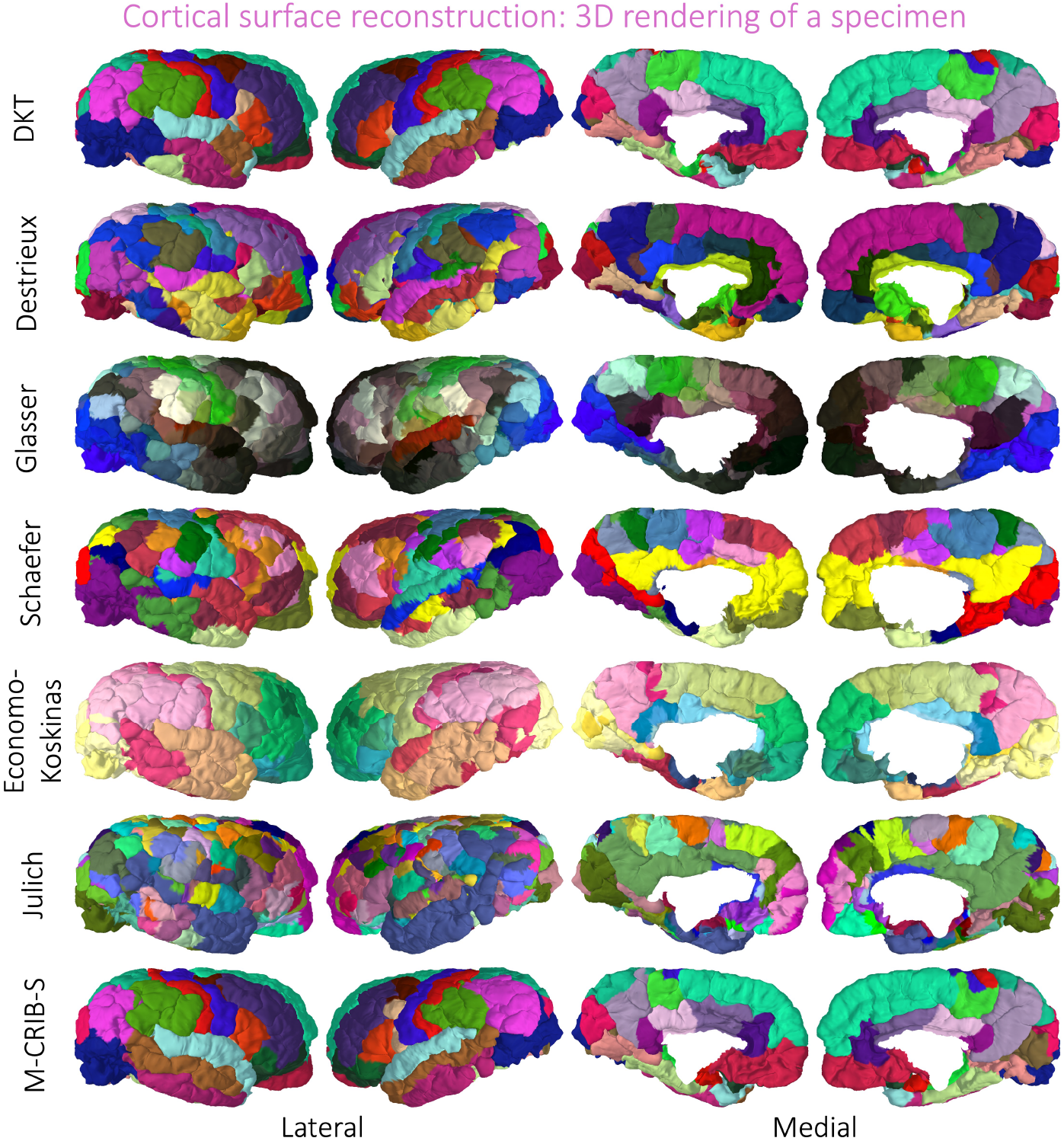
Postmortem human infant MRI. Each row depicts the reconstructed cortical surfaces based on the different atlases for the first specimen in Figure 3 for both hemispheres.

**Fig 5.**
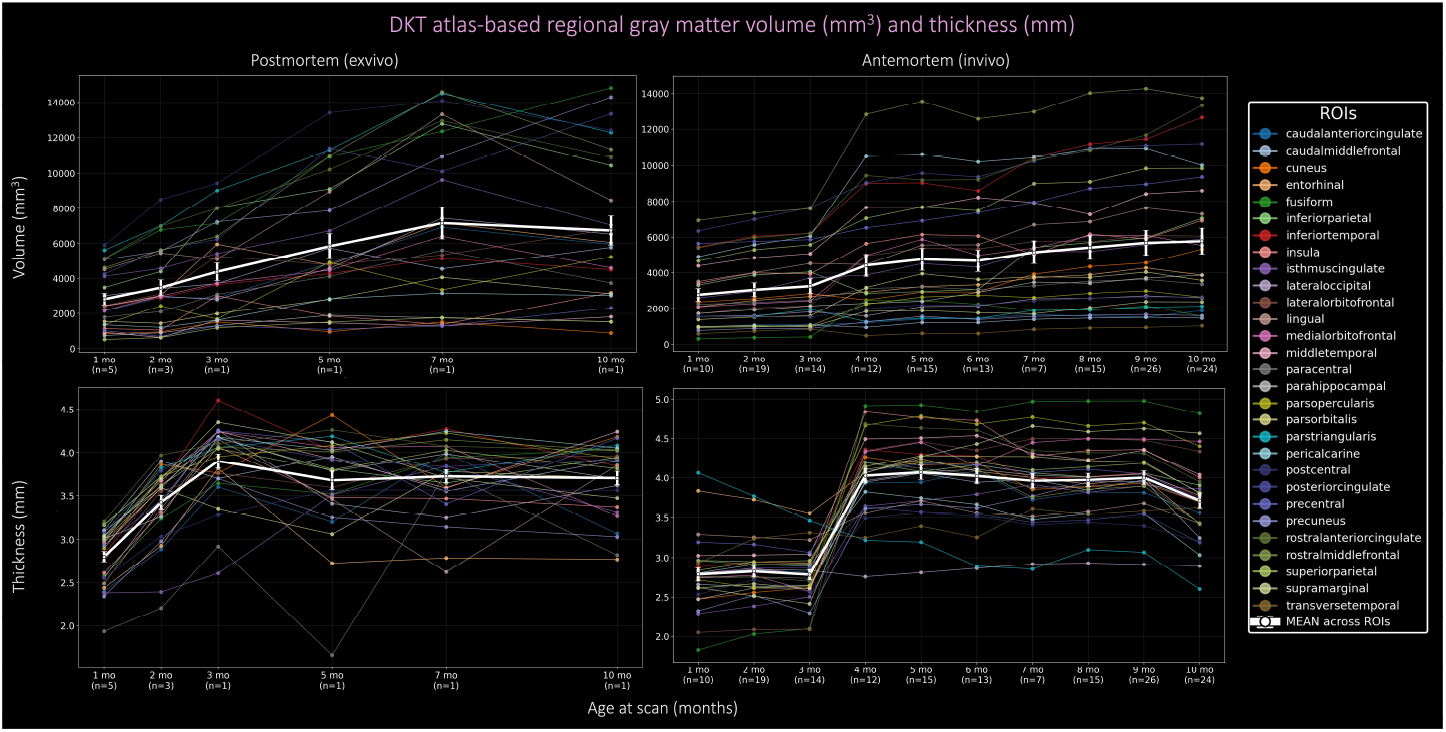
Postmortem brain morphometry. Desikan-Killiany-Tourville (DKT) atlas-based regional gray matter volumes and cortical thickness measurements are shown separately for postmortem MRI and *in vivo* BCP data from 0–10 months, yielding four plots: postmortem volume, postmortem thickness, BCP volume, and BCP thickness. Within each plot, colored curves represent individual DKT regions, and the bold white curve represents the mean across regions. Colors follow the standard FreeSurfer labeling scheme for the DKT cortical regions.

## 4 Discussion

We present a unified segmentation-to-surface reconstruction framework for highresolution multi-sequence postmortem infant MRI. The framework addresses a major gap in postmortem developmental neuroimaging by enabling automated tissue segmentation, cortical surface reconstruction, anatomical parcellation, and quantitative morphometric analysis in native postmortem space. Benchmarking against contrast-agnostic and foundation segmentation models demonstrates that existing general-purpose methods remain challenged by the unique combination of postmortem tissue preparation, infant neuroanatomy, incomplete myelination, altered tissue contrast, tissue deformation, and heterogeneous imaging sequences and acquisition protocols encountered in postmortem infant MRI. These results further suggest that matching infant anatomy alone is insufficient without accounting for postmortem-specific imaging characteristics.

An important contribution of the framework is its use of segmentation-derived anatomical boundaries to stabilize pial-surface placement when image-intensity gradients are unreliable. Robust cortical reconstruction and agreement of regional volumes and cortical thickness measurements with *in vivo* normative developmental trajectories support the anatomical plausibility and morphometric reliability of the pipeline. However, differences in tissue state, cohort composition, and processing pipelines preclude interpreting this comparison as direct morphometric validation.

The present study is limited by its small, single-site cohort, the absence of an independent external test set, the propagation of FLASH-based annotations to the other image types, and the lack of direct ground-truth surface measurements. External validation will therefore be necessary to establish generalization and surface accuracy across independent postmortem infant datasets. Future work will prioritize postmortem-specific synthetic data generation and foundation models trained on larger, more diverse postmortem infant datasets to further improve generalization across imaging protocols, tissue preparation conditions, and developmental stages.

## Acknowledgements

This work was supported by the National Institutes of Health (NIH) under grant R01HD102616.

## References

1. Olson, Halie A., et al. “Measuring and interpreting individual differences in fetal, infant, and toddler neurodevelopment.” Developmental Cognitive Neuroscience 73 (2025): 101539.

2. Chen, Liangjun, et al. “Four-dimensional mapping of dynamic longitudinal brain subcortical development and early learning functions in infants.” Nature Communications 14.1 (2023): 3727.

3. Shi, Feng, et al. “Infant brain atlases from neonates to 1-and 2-year-olds.” PloS one 6.4 (2011): e18746.

4. Ahmad, Sahar, et al. “Multifaceted atlases of the human brain in its infancy.” Nature Methods 20.1 (2023): 55–64.

5. Alex, Ann M., et al. “A global multicohort study to map subcortical brain development and cognition in infancy and early childhood.” Nature neuroscience 27.1 (2024): 176–186.

6. Dubois, Jessica, et al. “MRI of the neonatal brain: a review of methodological challenges and neuroscientific advances.” Journal of Magnetic Resonance Imaging 53.5 (2021): 1318–1343.

7. Iglesias, Juan Eugenio, et al. “A computational atlas of the hippocampal formation using ex vivo, ultra-high resolution MRI: Application to adaptive segmentation of in vivo MRI.” Neuroimage 115 (2015): 117–137.

8. Edlow, Brian L., et al. “7 Tesla MRI of the ex vivo human brain at 100 micron resolution.” Scientific data 6.1 (2019): 244.

9. Ravikumar, Sadhana, et al. “Postmortem imaging reveals patterns of medial temporal lobe vulnerability to tau pathology in Alzheimer’s disease.” Nature Communications 15.1 (2024): 4803.

10. Griffiths, P. D., M. N. J. Paley, and E. H. Whitby. “Post-mortem MRI as an adjunct to fetal or neonatal autopsy.” The Lancet 365.9466 (2005): 1271–1273.

11. Licandro, Roxane, et al. “High Resolution Postmortem MRI Discovers Developing Structural Connectivity in the Human Ascending Arousal Network.” Human Brain Mapping 46.17 (2025): e70422.

12. Khandelwal, Pulkit, et al. “Postmortem brain MRI reveals differential associations of subcortical and limbic volumes with cortical thinning and neurodegenerative pathologies.” Alzheimer’s & Dementia 22.7 (2026): e71649.

13. Tisdall, M. Dylan, et al. “Ex vivo MRI and histopathology detect novel iron-rich cortical inflammation in frontotemporal lobar degeneration with tau versus TDP-43 pathology.” NeuroImage: Clinical 33 (2022): 102913.

14. Kotrotsou, Aikaterini, et al. “Ex vivo MR volumetry of human brain hemispheres.” Magnetic resonance in medicine 71.1 (2014): 364–374.

15. Fischl, Bruce. “FreeSurfer.” Neuroimage 62.2 (2012): 774–781.

16. Jenkinson, Mark, et al. “Fsl.” Neuroimage 62.2 (2012): 782–790.

17. Henschel, Leonie, et al. “Fastsurfer-a fast and accurate deep learning based neuroimaging pipeline.” NeuroImage 219 (2020): 117012.

18. Fritz, Francisco J., et al. “Longitudinal whole-human-brain quantitative MRI study on autolysis, fixation, rehydration, and shrinkage effects.” bioRxiv (2026): 2026–01.

19. Khandelwal, Pulkit, et al. “Surface-based parcellation and vertex-wise analysis of ultra high-resolution ex vivo 7 tesla MRI in Alzheimer’s disease and related dementias.” International Workshop on Machine Learning in Clinical Neuroimaging. Cham: Springer Nature Switzerland, 2024.

20. Khandelwal, Pulkit, et al. “Automated deep learning segmentation of high-resolution 7 tesla postmortem MRI for quantitative analysis of structure-pathology correlations in neurodegenerative diseases.” Imaging Neuroscience 2 (2024): imag–2.

21. Casamitjana, Adrià, et al. “A probabilistic histological atlas of the human brain for MRI segmentation.” Nature 648.8094 (2025): 678–685.

22. Puonti, Oula, et al. “Fast segmentation with the NextBrain histological atlas.” Imaging Neuroscience (2026).

23. Chin, Jenna, et al. “Deep learning in fetal, infant, and toddler neuroimaging research.” Developmental Cognitive Neuroscience (2026): 101680.

24. Billot, Benjamin, et al. “SynthSeg: Segmentation of brain MRI scans of any contrast and resolution without retraining.” Medical image analysis 86 (2023): 102789.

25. Liu, Peirong, et al. “A modality-agnostic multi-task foundation model for human brain imaging.” arXiv preprint arXiv:2509.00549 (2025).

26. Shang, Ziyao, et al. “Learning strategies for contrast-agnostic segmentation via SynthSeg for infant MRI data.” International conference on medical imaging with deep learning. PMLR, 2022.

27. Hoffmann, Malte, Lilla Zöllei, and Adrian V. Dalca. “Deep infant brain segmentation from multi-contrast MRI.” arXiv preprint arXiv:2512.05114 (2025).

28. Zöllei, Lilla, et al. “Infant FreeSurfer: An automated segmentation and surface extraction pipeline for T1-weighted neuroimaging data of infants 0–2 years.” Neuroimage 218 (2020): 116946.

29. Wang, Li, et al. “iBEAT V2. 0: a multisite-applicable, deep learning-based pipeline for infant cerebral cortical surface reconstruction.” Nature protocols 18.5 (2023): 1488–1509.

30. Deoni, Sean CL, Terry M. Peters, and Brian K. Rutt. “High-resolution T1 and T2 mapping of the brain in a clinically acceptable time with DESPOT1 and DESPOT2.” Magnetic Resonance in Medicine: An Official Journal of the International Society for Magnetic Resonance in Medicine 53.1 (2005): 237–241.

31. Fischl, Bruce, et al. “Sequence-independent segmentation of magnetic resonance images.” Neuroimage 23 (2004): S69–S84.

32. https://github.com/pyushkevich/greedy

33. Dale, Anders M., Bruce Fischl, and Martin I. Sereno. “Cortical surface-based analysis: I. Segmentation and surface reconstruction.” Neuroimage 9.2 (1999): 179–194.

34. Fischl, Bruce, Martin I. Sereno, and Anders M. Dale. “Cortical surface-based analysis: II: inflation, flattening, and a surface-based coordinate system.” Neuroimage 9.2 (1999): 195–207.

35. Ségonne, Florent, Jenni Pacheco, and Bruce Fischl. “Geometrically accurate topology-correction of cortical surfaces using nonseparating loops.” IEEE transactions on medical imaging 26.4 (2007): 518–529.

36. Fischl, B., Sereno, M.I., Tootell, R.B. and Dale, A.M., 1999. High-resolution intersubject averaging and a coordinate system for the cortical surface. Human brain mapping, 8(4), pp.272–284.

37. Paszke, Adam, et al. “Pytorch: An imperative style, high-performance deep learning library.” Advances in neural information processing systems 32 (2019).

38. Isensee, Fabian, et al. “nnU-Net: a self-configuring method for deep learning-based biomedical image segmentation.” Nature methods 18.2 (2021): 203–211.

39. Isensee, Fabian, et al. “nnu-net revisited: A call for rigorous validation in 3d medical image segmentation.” International Conference on Medical Image Computing and Computer-Assisted Intervention. Cham: Springer Nature Switzerland, 2024.

40. Howell, Brittany R., et al. “The UNC/UMN Baby Connectome Project (BCP): An overview of the study design and protocol development.” NeuroImage 185 (2019): 891–905.

41. Adamson, Chris L., et al. “Parcellation of the neonatal cortex using Surface-based Melbourne Children’s Regional Infant Brain atlases (M-CRIB-S).” Scientific reports 10.1 (2020): 4359.

42. Klein, Arno, and Jason Tourville. “101 labeled brain images and a consistent human cortical labeling protocol.” Frontiers in neuroscience 6 (2012): 171.

43. Schaefer, Alexander, et al. “Local-global parcellation of the human cerebral cortex from intrinsic functional connectivity MRI.” Cerebral cortex 28.9 (2018): 3095–3114.

44. Amunts, Katrin, et al. “Julich-Brain: A 3D probabilistic atlas of the human brain’s cytoarchitecture.” Science 369.6506 (2020): 988–992.

45. Glasser, Matthew F., et al. “A multi-modal parcellation of human cerebral cortex.” Nature 536.7615 (2016): 171–178.

46. Scholtens, Lianne H., et al. “An mri von economo–koskinas atlas.” NeuroImage 170 (2018): 249–256.

